# Polygenic risk scores for major depressive disorder and neuroticism as predictors of antidepressant response: meta-analysis of three treatment cohorts

**DOI:** 10.1101/295717

**Authors:** Joey Ward, Nicholas Graham, Rona Strawbridge, Amy Ferguson, Gregory Jenkins, Wenan Chen, Mark Frye, Richard Weinshilboum, Rudolf Uher, Cathryn M. Lewis, Joanna Biernacka, Daniel J. Smith

**Affiliations:** University of Glasgow; Mayo Clinic; St. Jude Children’s Research Hospital; Dalhousie University; King’s College London

## Abstract

There are currently no reliable approaches for correctly identifying which patients with major depressive disorder (MDD) will respond well to antidepressant therapy. However, recent genetic advances suggest that Polygenic Risk Scores (PRS) could allow MDD patients to be stratified for antidepressant response. We used PRS for MDD and PRS for neuroticism as putative predictors of antidepressant response within three treatment cohorts: The Genome-based Therapeutic Drugs for Depression (GENDEP) cohort, and 2 sub-cohorts from the Pharmacogenomics Research Network Antidepressant Medication Pharmacogenomics Study PRGN-AMPS (total patient number = 783). Results across cohorts were combined via meta-analysis within a random effects model. Overall, PRS for MDD and neuroticism did not significantly predict antidepressant response but there was a consistent direction of effect, whereby greater genetic loading for both MDD (best MDD result, p < 5*10-5 MDD-PRS at 4 weeks, β = -0.019, S.E = 0.008, p = 0.01) and neuroticism (best neuroticism result, p < 0.1 neuroticism-PRS at 8 weeks, β = -0.017, S.E = 0.008, p = 0.03) were associated with less favourable response. We conclude that the PRS approach may offer some promise for treatment stratification in MDD and should now be assessed within larger clinical cohorts.

## Introduction

Major Depressive disorder (MDD) is a leading cause of disability worldwide (Ferrari *et al*, 2013). Antidepressants such as Selective Serotonin Reuptake Inhibitors (SSRIs) are first line treatments for MDD but up to one third of patients do not respond satisfactorily (Linde *et al*, 2015; Rush *et al*, 2006). There are currently no robust methods for predicting whether an individual patient will respond well to SSRIs and there is often a lag period of several weeks before clinical response, making decisions on switching to a different class of antidepressant difficult. Individual genetic variation may dictate likelihood of response to SSRIs (Peterson *et al*, 2017) and, as such, stratifying patients into sub-groups based on genetic profiles may allow for more efficient targeting of treatment.

Polygenic risk scoring (PRS) (Dudbridge, 2013) is a method which allows an individual’s genetic loading for a trait to be calculated using the output of genome-wide association study (GWAS) summary statistics. As current GWAS methodology does not capture the full extent of genetic effects on any given trait, a range of scores are created at different association p-value cut offs, allowing for the capture of more variance than that covered by only genome-wide significant loci.

It has been shown that a PRS can be of clinical use in predicting traits in independent samples. For example, for coronary heart disease, PRS improved the 10 year risk prediction in those over age 60 (Fernández-Ruiz, 2016). PRS approaches can also predict response to treatment, as demonstrated recently with an association between PRS for schizophrenia and less favourable response to lithium in bipolar disorder (International Consortium on Lithium, 2017). Here we test the hypothesis that PRS for MDD and PRS for neuroticism are associated with less favourable response to SSRIs, specifically citalopram and its active S-enantiomer escitalopram, in patients with MDD. Neuroticism is of particular interest in this regard because it is known to influence both serotonergic neurotransmission (Frokjaer *et al*, 2008) and response to antidepressants (Di Simplicio *et al*, 2014; Katon *et al*, 2010), and those with higher phenotypic neuroticism are less likely to respond as well to antidepressant therapy (Steffens *et al*, 2013).

## Methods

### Cohort descriptions, genotyping and imputation

Genome Based Therapeutic Drugs for Depression (GENDEP) is a cohort of 868 individuals, recruited from across Europe, treated with two classes of antidepressants: escitalopram (an SSRI) and nortriptyline (a tricyclic antidepressant). For the purposes of this study, only those patients in GENDEP treated with an SSRI were assessed (n = 267). Depressive symptoms were assessed on the Montgomery-Asberg Depression Rating Scale (MADRS), with measurements taken weekly for 12 weeks from baseline. Full genotyping and imputation methodology in GENDEP is described in previous reports (Uher *et al*, 2010).

The Pharmacogenomics Research Network Antidepressant Medication Pharmacogenomics Study (PGRN-AMPS) is a pharmacogenomics study of citalopram/escitalopram for treatment of MDD performed at the Mayo Clinic. An initial batch of 530 subjects (N=499 subjects of European ancestry that passed quality control) was genotyped for a pharmacogenomics GWAS of SSRIs (Ji *et al*, 2013). An additional 229 patients recruited in the PGRN-AMPS were subsequently genotyped for the International SSRI Pharmacogenomics Consortium (ISPC) GWAS (Biernacka *et al*, 2015). Depressive symptoms were assessed on the Hamilton Depression Rating Scale (HAMD). Full genotyping and imputation of these cohorts (here referred to as AMPS-1 and AMPS-2) have been described previously (Biernacka *et al*, 2015; Ji *et al*, 2013).

### Principal component generation and PRS construction

Principal genetic components were derived using PLINK. For all models the top 4 principal components were used as covariates in the model to account for hidden population structure. To ensure that an ethnically homogeneous sample was used in the AMPS-1 and AMPS-2 cohorts those whose Principal genetic components 1 to 4 were outside two standard deviations from the mean were excluded as outliers.

PRS were created using PLINK (Purcell *et al*, 2007) using outputs from the Smith et al. (2016) neuroticism GWAS (Smith *et al*, 2016) and the “probable MDD” phenotype of Howard et al (2017) MDD GWAS from UK Biobank (manuscript in press). SNPs were filtered by MAF < 0.01, HWE p<1*10^-6^ and imputation score < 0.8 before Linkage Disequilibrium (LD) clumping. SNPs were clumped using LD parameters of r^2^ >0.05 in a 500kb window. Selection of SNPs for each clump was based on which SNP had the lowest p value. In the event 2 SNPs in a clump had the same P value the SNP with the largest beta coefficient was selected. The scores generated were average scores with no-mean-imputation flag. Six profile scores were created for each trait using p value cut offs of p < 5*10^-8^, p < 5*10^-5^, p < 0.01, p < 0.05, p < 0.1 and p < 0.5. Risk scores were then standardised to mean =0, SD =1, and split into quintiles. Only those whose genetic loading was in the top and bottom quintile of each PRS were used in the analysis(Lewis and Vassos, 2017). For the GENDEP cohort the top and bottom quintile from each assessment centre was selected.

### Phenotype definition

For all three cohorts the primary outcome of interest was percentage change in depression score from baseline at four weeks. This was calculated by subtracting the score at four weeks from baseline, and dividing this difference by the score at baseline. A secondary outcome at eight weeks was also assessed, calculated using the same method. To be included in the analysis, an individual had to have a score recorded at baseline, four weeks and eight weeks.

### Statistical modelling

Modelling was performed in R using the lm function. All models were adjusted for age, sex and the first 4 principal components. The GENDEP models were additionally adjusted for recruitment centre which was treated as a factor variable. The r^2^ for the PRS term of the model was derived using the methodology described in Selzam et al(Selzam *et al*, 2016). Due to the results being largely null we did not perform any correction for multiple testing.

## Results

Demographic and clinical characteristics of the three cohorts (GENDEP, AMPS-1, and AMPS-2) are presented in table 1.

**Table 1.** Demographic and clinical characteristics.

| Cohort | Total N | N used in regressions | N female (%) | Age, mean(SD) | Baseline score, mean (SD) | 4 week score, mean (SD) | 8 week score, mean (SD) |
| --- | --- | --- | --- | --- | --- | --- | --- |
| AMPS-1 | 363 | 357 | 229 (64.1) | 40.9 (13.5) | 22 (4.88) | 11.9 (6.7) | 8.83 (5.92) |
| AMPS-2 | 145 | 138 | 85 (61.6) | 40.1 (13.6) | 21.2 (5.14) | 12 (5.84) | 9.14 (6.41) |
| GENDEP | 275 | 265 | 170 (64.2) | 42.3 (11.8) | 28.3 (6.16) | 18.7 (8.2) | 14.2 (8.89) |

\*score rating is HAMD for AMPS-1 and AMPS-2 and MADRS for GENDEP.

### Individual study analyses

The results of all the individual study analyses can be found in table s1–s3. Two of the models returned nominally significant results, both of which were in the AMPS-2 cohort (table 2). They were neuroticism p < 0.5 PRS at four weeks (β = -0.04, p=0.02) and neuroticism p < 0.5 at eight weeks (β = -0.039, p=0.03). Of particular note is the r^2^ of the PRS term of the significant models which accounts for approximately 10% of the variance. Note, however, that these results would not pass correction for multiple testing.

**Table 2.** Nominally significant individual PRS models (AMSP-2 cohort).

| **predictor** | **Time point**<br>**(weeks)** | **p** | **Beta** | **SE** | **T Test stat** | **r<sup>2</sup>** |
| --- | --- | --- | --- | --- | --- | --- |
| Neuroticism p<0.5 | 4 | 0.019 | -0.044 | 0.018 | -2.42 | 0.1 |
| Neuroticism p<0.5 | 8 | 0.029 | -0.039 | 0.017 | -2.26 | 0.08 |

Although we were unable to reject the null hypothesis in the rest of the models, a clear majority (56 of 72 models) identified beta coefficients in the same direction of effect (greater loading for MDD or neuroticism associated with a smaller percentage drop in depression score). Of the 16 positive beta coefficient models, ten were from GENDEP MDD PRS models, three were from GENDEP neuroticism PRS model, two were from AMPS-1 neuroticism PRS models and one was an AMPS-2 MDD PRS models (supplementary tables s1–s3).

### Meta-analysis

Two of the 24 meta-analyses were nominally significant: MDD p < 5*10^-5^ PRS at four weeks (β =-0.02, p=0.009, I^2^ = 0); and neuroticism p<0.1 PRS at eight weeks (β =-0.017, p=0.03, I^2^ = 0) (figure 1). Neither of these results would survive correction for multiple testing. The direction of effect in all of the meta-analyses was negative (greater genetic loading for MDD and neuroticism associated with a smaller percentage drop in depression score at both four and eight weeks; table S4. The forest plots of all other meta-analyses are provided as supplementary material.

**Figure 1.**
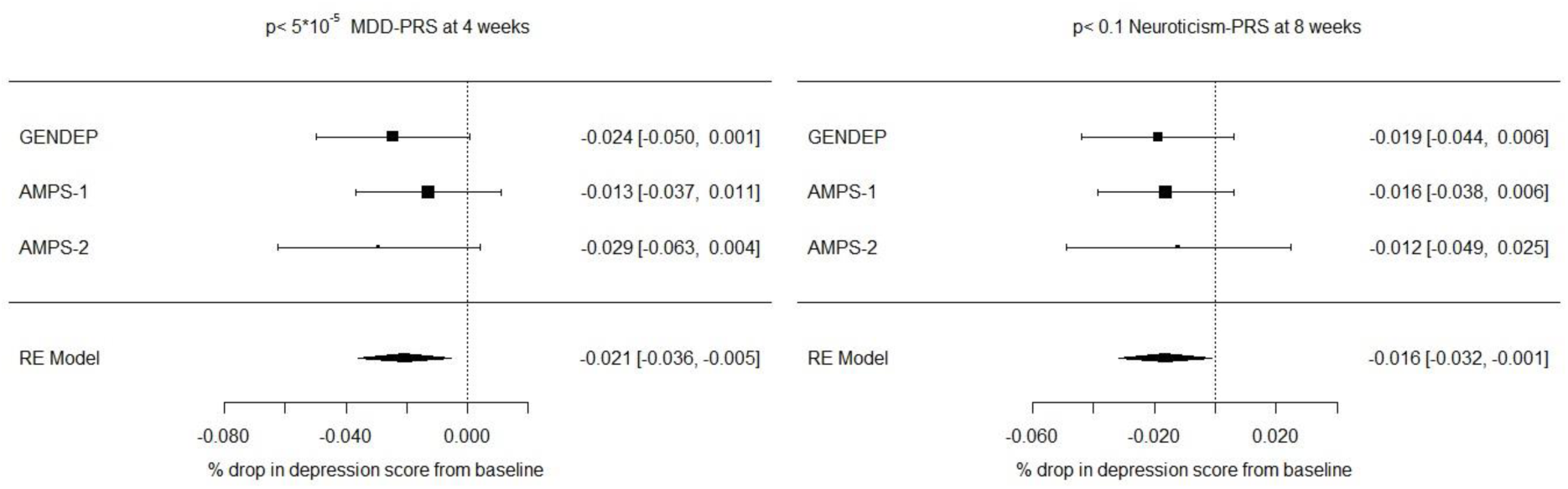
Forrest plot of nominally significant meta-analyses.

## Discussion

Our goal was to assess the extent to which PRS for MDD and PRS for neuroticism were associated with response to SSRIs in patients with MDD. Although most of the findings were null, there was a direction of effect where higher PRS for MDD and higher PRS for neuroticism were associated with less favourable response to SSRIs. It is likely that our analyses were under-powered – replication in larger datasets will therefore be of interest. We estimate that a training sample of approximately 10,000 and a target sample of 5,000 individuals would give 60% power in a PRS of 100,000 SNPs that explain 10% of the variance in the training sample (Palla and Dudbridge, 2015). For the two AMPS-2 nominally significant results the r^2^ values explaining approximately 10% of the variance, suggesting that these PRSs could potentially be useful clinically.

This work diverges from previous analyses in these cohorts which have focused on GWAS and candidate gene analyses to identify genetic loci that associate with antidepressant response. It is possible that the use of PRS is advantageous for clinical use over these methods as it allows for a whole-genome approach instead of focusing on specific SNPs, genes or regions. An individual’s response to antidepressants is likely to be influenced by many genetic factors and, as such, candidate gene methodologies will fail to capture polygenic influences. An additional strength of this work is that all three cohorts systematically assessed treatment response at comparable time-points and in the context of the use of the same class of antidepressants, namely SSRIs.

## Limitations

Apart from the issue of low power, our methodology was one in which only the extreme ends of genetic loadings were considered. This makes it difficult to translate the findings into a general population setting and routine clinical practice. Further work is needed to assess genetic loadings for MDD and neuroticism within the general population and how these relate to the clinical cohorts described here. The use of different depression rating scales between GENDEP and the AMPS- 1/AMPS-2 may have had some impact on the results as they may have captured different aspects of the depressive phenotype and symptom changes induced by antidepressants. However, I^2^ was low in the meta-analyses that achieved nominal significance. Using a consistent depression rating in future would aid in keeping heterogeneity consistently low.

Another limitation was in the estimation of LD blocks in the GENDEP cohort. Due to the cohort being composed of individuals across Europe, treating the group as a whole for estimating which SNPs are in LD may have led to inaccuracies. This could explain why many of coefficients in the GENDEP models showed as positive correlation unlike the models from AMPS-1 and AMPS-2. Principal component analysis of treatment centres showed overlapping clusters but they were not distinct enough to warrant calculating LD in each centre separately. Further work in this area should capture more detail on ethnicity and ancestral background, to allow for more robust determination of LD clumps and more informed decisions on the most appropriate inclusion criteria.

Finally, the result may have been impeded by the use of a single PRS predictor. Recent research has shown that the use of multiple scores covering a variety genetic loadings can explain significantly more variance that that of a single score (Krapohl *et al*, 2017). As such, for outcomes as complex as antidepressant response may prove more fruitful.

## Conclusion

Stratified medicine in psychiatry is still in its infancy. Genotyping is not currently routine practice in clinical settings and the use of PRS to guide the use of SSRIs in MDD remains a long-term goal.

However, with increasingly large and well-phenotyped cohorts available for analysis and more powerful GWAS outputs being produced, we tentatively conclude that more targeted prescribing of anti-depressants in MDD based on genetic profiles is a realistic prospect for the future.

## Funding and Disclosure

The main body of work for the analysis was performed by Joey Ward who is funded by the Royal college of Physicians of Edinburgh through the JMAS Sims fellowship for his project “*Towards precision medicine for depressive disorders”*.

## Conflicts of interest

none

## Supplementary tables

**Table S1.** Results of the individual regressions in the AMPS-1 cohort

| Time point (week) | predictor | P | Beta | SE | T stat | r<sup>2</sup> |
| --- | --- | --- | --- | --- | --- | --- |
| 4 | mdd $p < 5 \times 10^{-8}$ | 0.24 | -0.014 | 0.0118 | -1.17 | 0.01 |
| | mdd $p < 5 \times 10^{-5}$ | 0.03 | -0.013 | 0.0122 | -1.05 | 0.008 |
| | mdd $p < 0.01$ | 0.57 | -0.007 | 0.0122 | -0.571 | 0.002 |
| | mdd $p < 0.05$ | 0.43 | -0.0096 | 0.012 | -0.799 | 0.005 |
| | mdd $p < 0.1$ | 0.40 | -0.011 | 0.0125 | -0.853 | 0.005 |
| | mdd $p < 0.5$ | 0.12 | -0.02 | 0.0127 | -1.58 | 0.017 |
| 8 | mdd $p < 5 \times 10^{-8}$ | 0.19 | -0.015 | 0.0116 | -1.31 | 0.012 |
| | mdd $p < 5 \times 10^{-5}$ | 0.18 | -0.017 | 0.0122 | -1.35 | 0.013 |
| | mdd $p < 0.01$ | 0.14 | -0.016 | 0.0107 | -1.48 | 0.015 |
| | mdd $p < 0.05$ | 0.49 | -0.0077 | 0.0111 | -0.695 | 0.003 |
| | mdd $p < 0.1$ | 0.47 | -0.0078 | 0.0108 | -0.72 | 0.004 |
| | mdd $p < 0.5$ | 0.11 | -0.018 | 0.0111 | -1.59 | 0.017 |
| 4 | neuroticism $p < 5 \times 10^{-8}$ | 0.77 | -0.0036 | 0.0125 | -0.288 | 0.0006 |
| | neuroticism $p < 5 \times 10^{-5}$ | 0.46 | -0.0095 | 0.0128 | -0.744 | 0.004 |
| | neuroticism $p < 0.01$ | 0.85 | -0.0024 | 0.0126 | -0.192 | 0.0003 |
| | neuroticism $p < 0.05$ | 0.39 | -0.011 | 0.0124 | -0.87 | 0.005 |
| | neuroticism $p < 0.1$ | 0.70 | -0.0048 | 0.0124 | -0.385 | 0.001 |
| | neuroticism $p < 0.5$ | 0.44 | 0.0091 | 0.0118 | 0.769 | 0.004 |
| 8 | neuroticism $p < 5 \times 10^{-8}$ | 0.88 | -0.0017 | 0.0115 | -0.146 | 0.0002 |
| | neuroticism $p < 5 \times 10^{-5}$ | 0.87 | -0.0018 | 0.0115 | -0.159 | 0.0002 |
| | neuroticism $p < 0.01$ | 0.68 | -0.0049 | 0.0117 | -0.42 | 0.001 |
| | neuroticism $p < 0.05$ | 0.17 | -0.016 | 0.0116 | -1.39 | 0.013 |
| | neuroticism $p < 0.1$ | 0.15 | -0.016 | 0.0113 | -1.43 | 0.015 |
| | neuroticism $p < 0.5$ | 0.95 | 0.0007 | 0.0105 | 0.0661 | 0.00003 |

**Table S2.** Results of the induvial regressions in the AMPS-2 cohort

| Time point<br>(week) | predictor | P | Beta | SE | T stat | r<sup>2</sup> |
| --- | --- | --- | --- | --- | --- | --- |
| 4 | mdd $p < 5 \times 10^{-8}$ | 0.80 | 0.004 | 0.02 | 0.25 | 0.001 |
| | mdd $p < 5 \times 10^{-5}$ | 0.090 | -0.03 | 0.02 | -1.73 | 0.039 |
| | mdd $p < 0.01$ | 0.06 | -0.03 | 0.02 | -1.93 | 0.066 |
| | mdd $p < 0.05$ | 0.32 | -0.01 | 0.01 | -1.01 | 0.018 |
| | mdd $p < 0.1$ | 0.60 | -0.01 | 0.02 | -0.52 | 0.005 |
| | mdd $p < 0.5$ | 0.58 | -0.01 | 0.02 | -0.55 | 0.005 |
| 8 | mdd $p < 5 \times 10^{-8}$ | 0.78 | -0.01 | 0.02 | -0.28 | 0.001 |
| | mdd $p < 5 \times 10^{-5}$ | 0.25 | -0.02 | 0.02 | -1.18 | 0.024 |
| | mdd $p < 0.01$ | 0.78 | -0.01 | 0.02 | -0.28 | 0.001 |
| | mdd $p < 0.05$ | 0.23 | -0.02 | 0.02 | -1.22 | 0.027 |
| | mdd $p < 0.1$ | 0.40 | -0.02 | 0.02 | -0.84 | 0.011 |
| | mdd $p < 0.5$ | 0.15 | -0.03 | 0.02 | -1.45 | 0.037 |
| 4 | neuroticism $p < 5 \times 10^{-8}$ | 0.20 | -0.03 | 0.02 | -1.3 | 0.03 |
| | neuroticism $p < 5 \times 10^{-5}$ | 0.37 | -0.01 | 0.02 | -0.91 | 0.01 |
| | neuroticism $p < 0.01$ | 0.079 | -0.03 | 0.02 | -1.79 | 0.06 |
| | neuroticism $p < 0.05$ | 0.14 | -0.03 | 0.02 | -1.49 | 0.04 |
| | neuroticism $p < 0.1$ | 0.28 | -0.02 | 0.02 | -1.1 | 0.02 |
| | neuroticism $p < 0.5$ | 0.019 | -0.04 | 0.02 | -2.42 | 0.099 |
| 8 | neuroticism $p < 5 \times 10^{-8}$ | 0.32 | -0.02 | 0.02 | -1 | 0.016 |
| | neuroticism $p < 5 \times 10^{-5}$ | 0.28 | -0.02 | 0.02 | -1.09 | 0.019 |
| | neuroticism $p < 0.01$ | 0.07 | -0.03 | 0.02 | -1.82 | 0.05 |
| | neuroticism $p < 0.05$ | 0.62 | -0.01 | 0.02 | -0.49 | 0.004 |
| | neuroticism $p < 0.1$ | 0.52 | -0.01 | 0.02 | -0.64 | 0.007 |
| | neuroticism $p < 0.5$ | 0.02 | -0.04 | 0.02 | -2.26 | 0.08 |

**Table S3.** Results of the induvial regressions in the GENDEP cohort

| Time point (week) | predictor | P | Beta | SE | T stat | r<sup>2</sup> |
| --- | --- | --- | --- | --- | --- | --- |
| 4 | mdd $p < 5 \times 10^{-8}$ | 0.77 | 0.004 | 0.013 | 0.3 | 0.0008 |
| | mdd $p < 5 \times 10^{-5}$ | 0.06 | -0.02 | 0.013 | -1.9 | 0.026 |
| | mdd $p < 0.01$ | 0.77 | 0.003 | 0.012 | 0.3 | 0.0007 |
| | mdd $p < 0.05$ | 0.51 | 0.008 | 0.012 | 0.66 | 0.003 |
| | mdd $p < 0.1$ | 0.62 | 0.006 | 0.013 | 0.5 | 0.002 |
| | mdd $p < 0.5$ | 0.52 | 0.009 | 0.013 | 0.64 | 0.003 |
| 8 | mdd $p < 5 \times 10^{-8}$ | 0.18 | 0.019 | 0.014 | 1.35 | 0.02 |
| | mdd $p < 5 \times 10^{-5}$ | 0.49 | -0.012 | 0.017 | -0.69 | 0.004 |
| | mdd $p < 0.01$ | 0.73 | 0.006 | 0.017 | 0.35 | 0.001 |
| | mdd $p < 0.05$ | 0.28 | 0.018 | 0.017 | 1.08 | 0.009 |
| | mdd $p < 0.1$ | 0.43 | 0.013 | 0.016 | 0.8 | 0.005 |
| | mdd $p < 0.5$ | 0.2 | 0.02 | 0.016 | 1.29 | 0.01 |
| 4 | neuroticism $p < 5 \times 10^{-8}$ | 0.71 | -0.004 | 0.011 | -0.37 | 0.001 |
| | neuroticism $p < 5 \times 10^{-5}$ | 0.62 | -0.006 | 0.012 | -0.5 | 0.002 |
| | neuroticism $p < 0.01$ | 0.29 | -0.013 | 0.012 | -1.06 | 0.009 |
| | neuroticism $p < 0.05$ | 0.97 | -0.001 | 0.012 | -0.04 | 0.00001 |
| | neuroticism $p < 0.1$ | 0.33 | -0.01 | 0.012 | -0.97 | 0.008 |
| | neuroticism $p < 0.5$ | 0.61 | -0.006 | 0.012 | -0.5 | 0.002 |
| 8 | neuroticism $p < 5 \times 10^{-8}$ | 0.99 | 0.0001 | 0.012 | 0.005 | 0.0000002 |
| | neuroticism $p < 5 \times 10^{-5}$ | 0.97 | 0.0005 | 0.013 | 0.04 | 0.00001 |
| | neuroticism $p < 0.01$ | 0.92 | 0.002 | 0.018 | 0.095 | 0.00008 |
| | neuroticism $p < 0.05$ | 0.36 | -0.012 | 0.013 | -0.91 | 0.007 |
| | neuroticism $p < 0.1$ | 0.15 | -0.019 | 0.013 | -1.47 | 0.02 |
| | neuroticism $p < 0.5$ | 0.43 | -0.01 | 0.013 | -0.79 | 0.005 |

**Table S4.** Results of the meta-analyses

| outcome | PRS | beta | SE | Z | P | ci.lb | ci.ub | I2 |
| --- | --- | --- | --- | --- | --- | --- | --- | --- |
| % drop in 4 weeks | MDD $p < 5*10^{-8}$ | -0.004 | 0.008 | -0.49 | 0.63 | -0.02 | 0.01 | 0 |
| | MDD $p < 5*10^{-5}$ | -0.021 | 0.008 | -2.63 | 0.01 | -0.04 | -0.01 | 0 |
| | MDD $p < 0.01$ | -0.009 | 0.009 | -1 | 0.32 | -0.03 | 0.01 | 28.2 |
| | MDD $p < 0.05$ | -0.004 | 0.007 | -0.61 | 0.54 | -0.02 | 0.01 | 0 |
| | MDD $p < 0.1$ | -0.004 | 0.008 | -0.48 | 0.63 | -0.02 | 0.01 | 0 |
| | MDD $p < 0.5$ | -0.007 | 0.009 | -0.77 | 0.44 | -0.02 | 0.01 | 21.6 |
| % drop in 8 weeks | MDD $p < 5*10^{-8}$ | -0.001 | 0.012 | -0.09 | 0.93 | -0.02 | 0.02 | 45.3 |
| | MDD $p < 5*10^{-5}$ | -0.017 | 0.009 | -1.86 | 0.06 | -0.03 | 0.00 | 0 |
| | MDD $p < 0.01$ | -0.009 | 0.008 | -1.08 | 0.28 | -0.02 | 0.01 | 0 |
| | MDD $p < 0.05$ | -0.004 | 0.010 | -0.36 | 0.72 | -0.02 | 0.02 | 23.9 |
| | MDD $p < 0.1$ | -0.004 | 0.008 | -0.50 | 0.62 | -0.02 | 0.01 | 0 |
| | MDD $p < 0.5$ | -0.008 | 0.014 | -0.58 | 0.57 | -0.04 | 0.02 | 61.5 |
| % drop in 4 weeks | N $p < 5*10^{-8}$ | -0.007 | 0.008 | -0.94 | 0.35 | -0.02 | 0.01 | 0 |
| | N $p < 5*10^{-5}$ | -0.009 | 0.008 | -1.17 | 0.24 | -0.02 | 0.01 | 0 |
| | N $p < 0.01$ | -0.012 | 0.008 | -1.56 | 0.12 | -0.03 | 0.00 | 3 |
| | N $p < 0.05$ | -0.009 | 0.008 | -1.17 | 0.24 | -0.02 | 0.01 | 0 |
| | N $p < 0.1$ | -0.010 | 0.008 | -1.33 | 0.18 | -0.03 | 0.00 | 0 |
| | N $p < 0.5$ | -0.011 | 0.015 | -0.75 | 0.45 | -0.04 | 0.02 | 70.4 |
| % drop in 8 weeks | N $p < 5*10^{-8}$ | -0.004 | 0.008 | -0.49 | 0.62 | -0.02 | 0.01 | 0 |
| | N $p < 5*10^{-5}$ | -0.004 | 0.008 | -0.55 | 0.58 | -0.02 | 0.01 | 0 |
| | N $p < 0.01$ | -0.010 | 0.009 | -1.09 | 0.27 | -0.03 | 0.01 | 2.3 |
| | N $p < 0.05$ | -0.013 | 0.008 | -1.70 | 0.09 | -0.03 | 0.00 | 0 |
| | N $p < 0.1$ | -0.017 | 0.008 | -2.13 | 0.03 | -0.03 | 0.00 | 0 |
| | N $p < 0.5$ | -0.013 | 0.011 | -1.20 | 0.23 | -0.03 | 0.01 | 48 |

## Supplementary Figures

**Figure S1.**
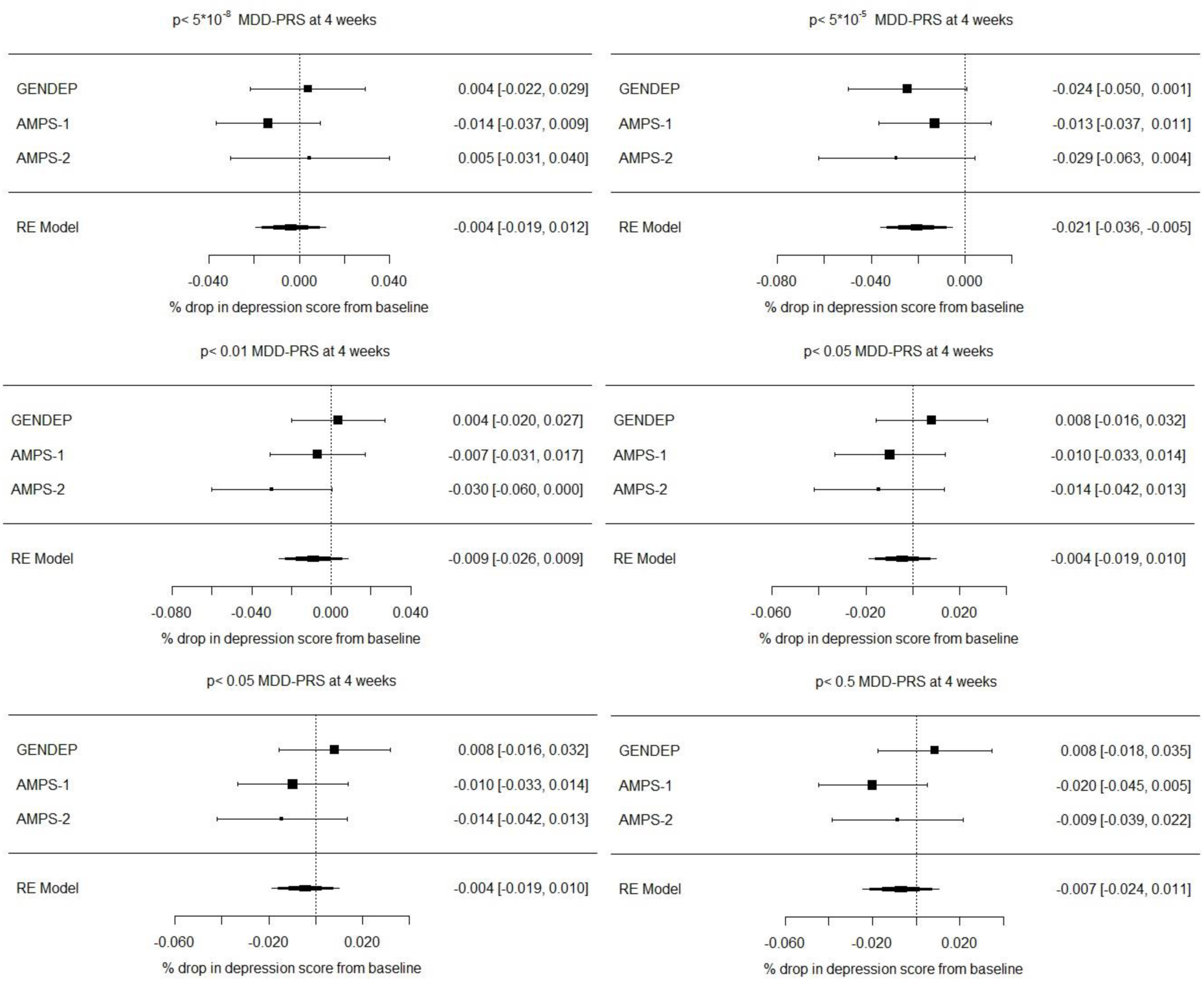
MDD PRS meta-analysis results at 4 weeks

**Figure S2.**
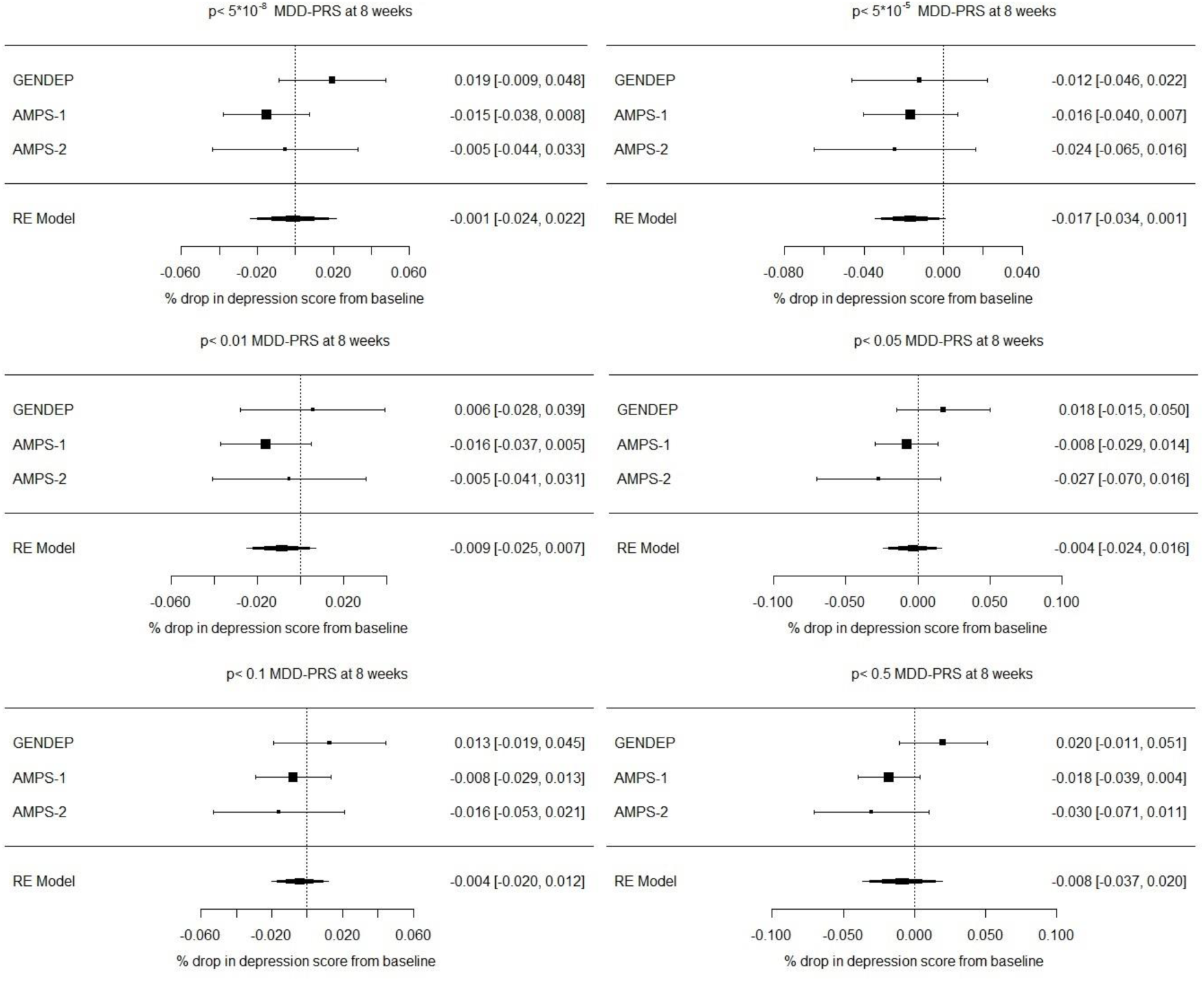
MDD PRS meta-analysis results at 8 weeks

**Figure S3.**
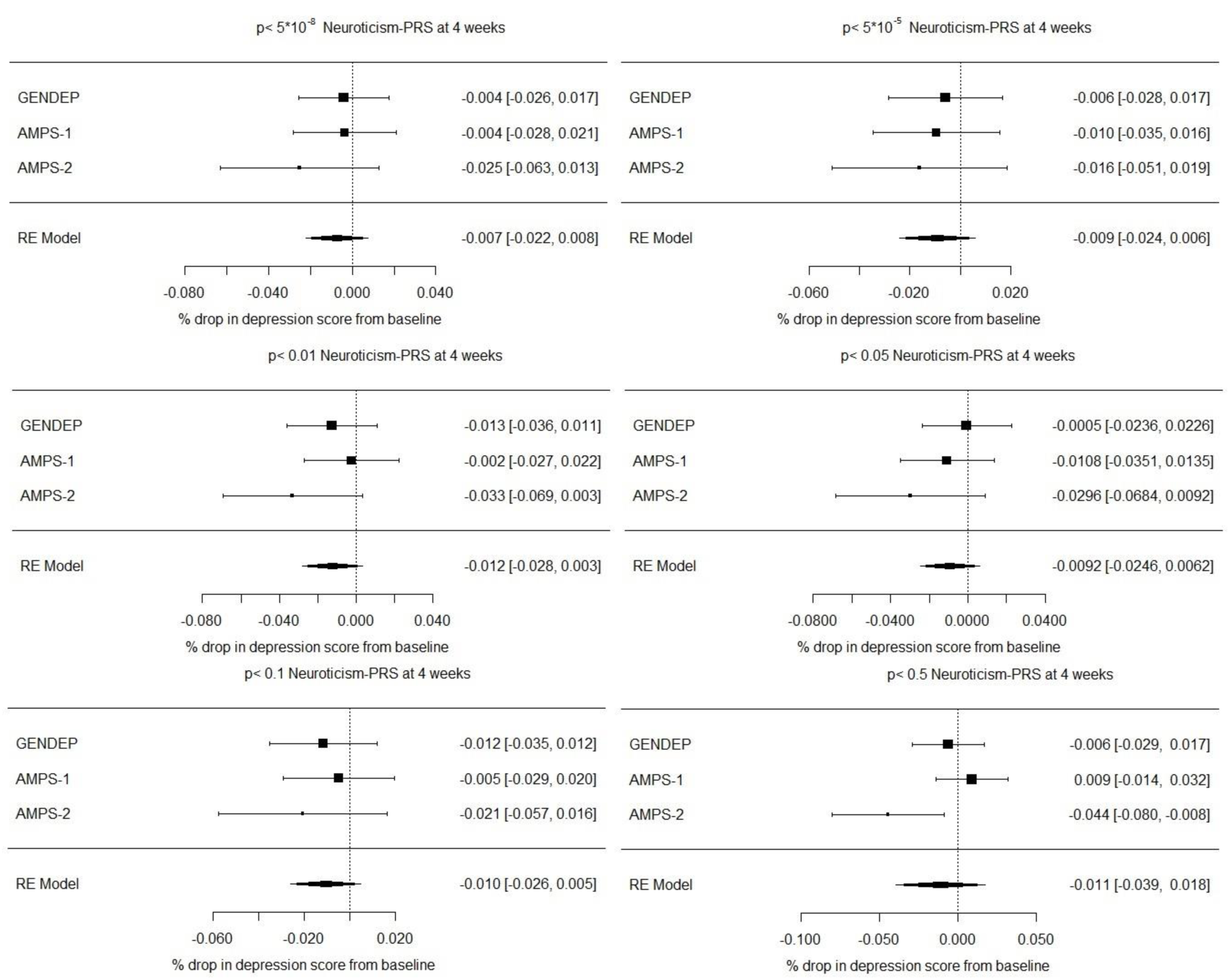
Neuroticism PRS meta-analysis results at 4 weeks

**Figure S4.**
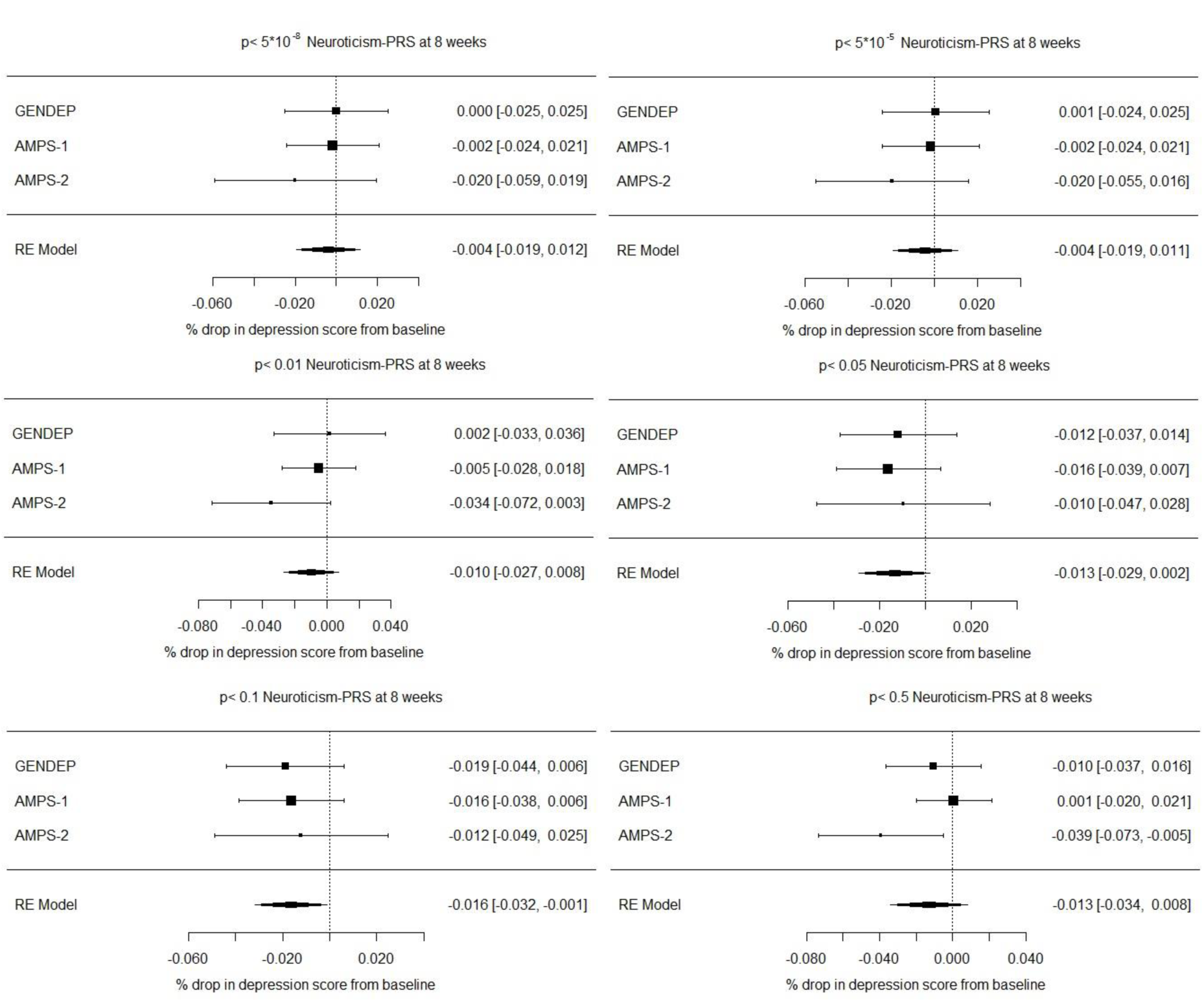
Neuroticism PRS meta-analysis results at 8 weeks

